# Bimodal Whole-Mount Imaging Of Tendon Using Confocal Microscopy And X-Ray Micro-Computed Tomography

**DOI:** 10.1101/2020.04.08.031450

**Authors:** Neil Marr, Mark Hopkinson, Andrew P. Hibbert, Andrew A. Pitsillides, Chavaunne T. Thorpe

**Author notes:** **CORRESPONDING AUTHOR:** Chavaunne Thorpe.

## Abstract

**BACKGROUND:** 3-dimensional imaging modalities for optically dense connective tissues such as tendons are limited and typically have a single imaging methodological endpoint. Here, we have developed a bimodal procedure that utilises fluorescence-based confocal microscopy and x-ray micro-computed tomography for the imaging of adult tendons to visualise and analyse extracellular sub-structure and cellular composition in small and large animal species.

**RESULTS:** Using fluorescent immunolabelling and optical clearing, we visualised the expression of the basement membrane protein laminin-α4 in 3D throughout whole rat Achilles tendons and equine superficial digital flexor tendon 5 mm segments. This revealed a complex network of laminin-α4 within the tendon core that predominantly localises to the interfascicular matrix compartment. Furthermore, we implemented a chemical drying process capable of creating contrast densities enabling visualisation and quantification of both fascicular and interfascicular matrix volume and thickness by x-ray micro-computed tomography. We also demonstrated that both modalities can be combined using reverse clarification of fluorescently labelled tissues prior to chemical drying to enable bimodal imaging of a single sample.

**CONCLUSIONS:** Whole-mount imaging of tendon allowed us to identify the presence of an extensive network of laminin-α4 within tendon, the complexity of which cannot be appreciated using traditional 2D imaging techniques. Creating contrast for x-ray micro-computed tomography imaging of tendon using chemical drying is not only simple and rapid, but also markedly improves on previously published methods. Combining these methods provides the ability to gain spatio-temporal information and quantify tendon substructures to elucidate the relationship between morphology and function.

## INTRODUCTION

Advances in 3-dimensional (3D) imaging of dense connective tissues such as tendons are essential for the investigation of normal tissue structure as well as musculoskeletal diseases in pre-clinical models and clinical samples. Recent developments in 3D microscopy and scanning techniques have permitted imaging of cells and structures of calcified tissues, whole embryos, and organisms, using methods including phase-contrast x-ray micro-computed tomography (μ-CT), optical projection tomography and label-free detection methods [1–3]. However, 3D imaging by fluorescent methods remains a challenge for adult tissues such as cartilage, ligaments and tendons, as their opacity and dense matrix composition renders deep imaging of whole connective tissues difficult. Paradoxically, x-ray micro-computed tomography (μ-CT) of non-calcified tissues is technically difficult due to their lower x-ray attenuation compared to mineralised tissues such as bone [4]. Hence, there is a demand for imaging modalities that can be used to study the gross structure of connective tissues as well as the spatial organisation of extracellular matrix (ECM) and its inter-relationships with resident cell populations.

Until recently, imaging techniques to investigate both structural and cellular elements of dense collagenous tissues such as adult tendon have been limited to conventional 2D methods. These only allow appreciation of tissue structure in a single plane or require extensive reconstruction [5], and are time-consuming, labour-intensive, and destructive, often creating artefacts within tissue [6]. Recent advances in optical clearing agents have provided scope to clarify tissues, either by dehydration, delipidation, matching tissue refractive index or a combination of each, to allow 3D visualisation of ECM organisation and cell populations in both mineralised and non-mineralised tissues [7–11]. A plethora of clearing agents are now commercially available, with a number of studies describing their effectiveness for fluorescent imaging of connective tissues with varying degrees of success [12–15]. In addition, reversing optical clarification of collagenous structures is possible with a variety of aqueous compounds, such as rehydration by saline-based solutions of glycerol or benzyl benzoate based clearing agents [13, 16]. Visikol® HISTO™ is a clearing agent reversible by ethanol which has only minor effects on tissue structure [17], with recent studies able to reverse tissue clearing for histological imaging post-3D imaging [18, 19]. Therefore, the reversibility of clarification agents introduces a new potential to better integrate different imaging modalities to resolve tissue structure and cell-ECM relationships. To the authors’ knowledge, no study to date has attempted to establish bimodal procedures to image fluorescently labelled soft tissues in 3D and apply a distinct modality, such as μ-CT, to assess gross structural parameters quantitatively. Hence, post-clarification imaging by μ-CT could provide a new avenue for soft tissue research to investigate structure-function relationships in conjunction with 3D immunolabelling approaches.

Contrast-enhanced 3D imaging in soft tissues remains difficult due to current strategies being limited by long perfusion times and an inability to fully resolve tissue structure due to insufficient contrast. Recent studies have described procedures to image soft tissues such as tendon with μ-CT using common aqueous contrast agents for such as iodine potassium iodide (I_2_KI; also referred to as Lugol’s solution) and phosphotungstic acid [20]. However, acidic agents such as phosphotungstic acid require high concentrations to create contrast which can erode samples and distort structure in soft tissues, whereas iodine-based agents require variable incubation times and may ultimately provide inadequate contrast to visualise and segment soft tissue sub-structures [20]. Hence, alternatives such as chemical and critical point drying should be considered, given both methods have been shown to generate excellent contrast in soft tissues [21–25]. Recent advances have exploited chemical drying by hexamethyldisilazane (HMDS) to generate contrast for 3D visualisation of soft tissue internal structure, although this technique has yet to be applied to tendon [26, 27].

Combining imaging that reveals internal structure of connective tissues with markers of cell:ECM specialisation in a single sample is an attractive possibility. In tendons, a number of fibrous proteins, proteoglycans, and glycoproteins that comprise basement membranes have been shown to localise to the interfascicular matrix (IFM), which surrounds tendon fascicles (reviewed in [28]). Basement membranes are highly specialised ECMs that interact with cell surface receptors, and are integral to progenitor cell niches, governing fate determination, structural integrity in musculoskeletal tissues (reviewed in [27] and [28]). Laminin is an essential basement membrane glycoprotein that interacts with cell adhesion molecules and cell surface receptors to regulate various cellular and molecular processes [30, 31]. In tendon, laminin and other basement membrane-associated proteins have been described previously [29, 30], with recent proteomic studies identifying numerous laminin subunits, including laminin α-4 (LAMA4), localised to the IFM [31]. However, the 3D organisation and composition of basement membrane proteins within the tendon core has yet to be described. Therefore, we utilised a novel cross-species marker of tendon basement membrane, laminin α-4 (LAMA4), combined with optical clearing techniques, to visualise 3D organisation of structural components in tendon.

Given the demand for 3D imaging modalities for dense connective tissues, we developed techniques for whole-mount confocal microscopy and x-ray micro-computed tomography to achieve 3D visualisation and analysis of adult tendons. Furthermore, we aimed to develop a method applicable to tissues from both small and large animal models, testing our method in a rat (*Rattus norvegicus*) and horse (*Equus caballus*) tendon. Next, we developed a workflow for bimodal single sample imaging, combining our confocal and μ-CT imaging methodologies to visualise cellular and structural properties of tendon in 3D.

## RESULTS

To develop a method that enables both fluorescent imaging of cell-ECM architecture and x-ray scanning of dense connective tissue structure, we combined multiple processes to create a workflow that integrates whole-mount immunolabelling, reversible optical clarification, and subsequent chemical drying to enable bimodal 3D imaging of tendon by confocal microscopy and μ-CT (**Figure 1**). We applied this technique to whole rat Achilles tendons and segments of equine superficial digital flexor tendons (SDFT).

**Figure 1.**
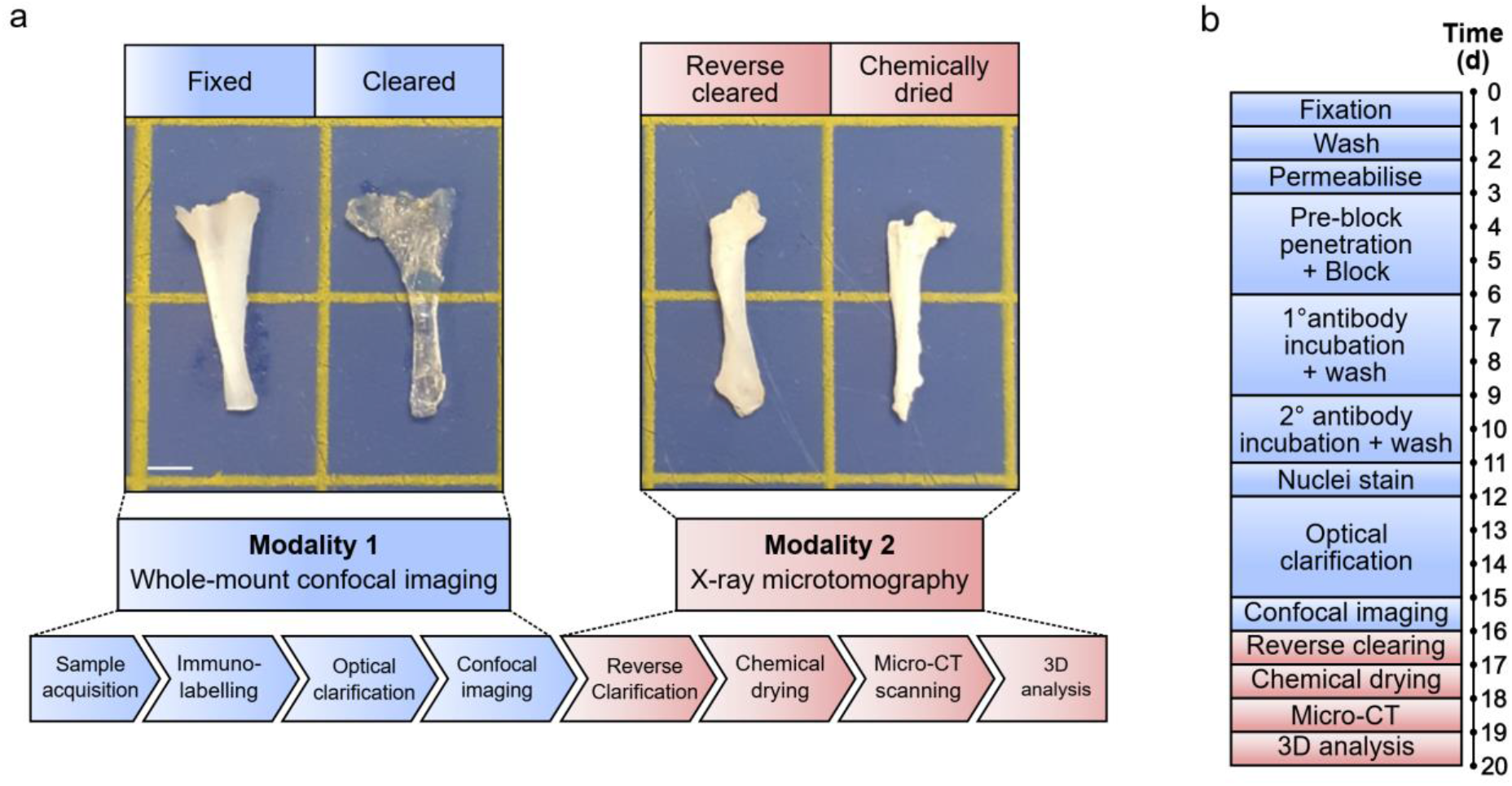
Visual workflow of bimodal tendon imaging by confocal microscopy and x-ray micro-computed tomography. (A) Schematic workflow of bimodal imaging of tendon with representative images of rat tendon after key processes in the protocol. Each modality can be performed independently of the other. Scale bar = 2.5 mm. (B) Visual timeline of typical bimodal 3D imaging modalities for whole rat Achilles and equine SDFT segment. Preparation for our optimised whole-mount confocal imaging takes a minimum of 16 days, whilst μ-CT modalities can be performed in 3 days following reverse clarification.

Prior to 3D imaging, appropriate antibody concentrations were identified and expression of LAMA4 was confirmed in both rat Achilles and equine SDFT using traditional 2D fluorescent immunolabelling of tendon sections (**Figure S1**). Thereafter, we performed tissue immunolabelling and optical clarification with Visikol HISTO™ solutions to visualise 3D organisation of LAMA4 in whole rat Achilles and in segments of equine SDFT using confocal microscopy using optimised protocols based on size-specific guidelines provided by Visikol™. As tissue thickness was similar between both equine SDFT segments and rat Achilles tendon, initial immunolabelling steps, including washing, permeabilisation, detergent washes and blocking were performed for identical periods of times. The main difference between protocols were antibody concentrations, which differed based on initial optimisations using thin tissue sections. The entire protocol, including nuclei counterstaining and dehydration steps prior to optical clarification requires a minimum of 12 days with no stopping points.

Based on our optimised protocol, clarification rendered both SDFT segments and whole rat Achilles transparent within 4.5 days. This was achieved with minimal clearing times (SDFT = 72 h; rat Achilles = 108 h) according to the manufacturer’s guidelines, and longer clarification steps can be used without negatively affecting labelling or tissue integrity. Using 3D projections of image stacks acquired by confocal imaging, we were able to visualise widespread vessel-like organisation of the structures labelled positively for basement membrane protein LAMA4 within the rat Achilles and equine SDFT (**Figure 2 & Figure 4**); information that cannot be gained from traditional 2D imaging. In the rat Achilles, LAMA4 expression was localised to the tendon surface and structures with a distribution likely to comprise the IFM (**Figure 2 & Figure 3**). In equine SDFT segments taken from the tendon core, the majority of LAMA4 labelling was seemingly consistent with IFM with some LAMA4 deposition in fascicles (**Figure 4**). Our method achieved imaging depths of at least 200 microns in equine SDFT and throughout an entire rat Achilles tendon (approximately 800 microns).

**Figure 2.**
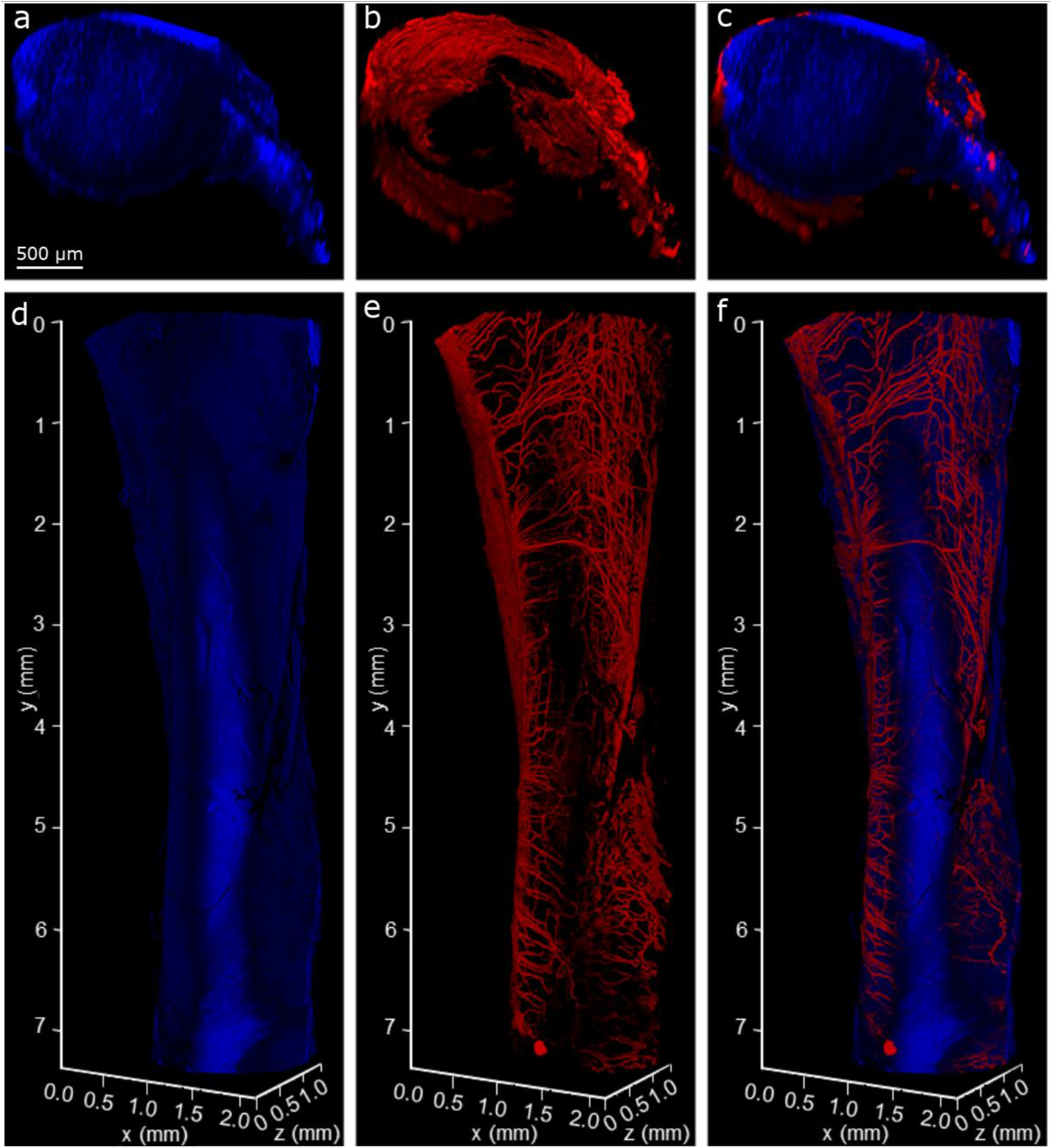
3D visualisation of whole-mount LAMA4-immunolabelled rat Achilles tendon. 3D reconstructions of a rat Achilles tendon showing nuclear (a,d,h) and LAMA4 (b,e,i) labelling, and with each channel overlaid to create a composite image (c,f,j). Transverse (xz) views of the 3D reconstructions (d-f) and tendon core with LAMA4-labelled IFM (h-j; magnification of white box) demonstrate signal present throughout the depth of the tissue. Longitudinal views (a-c) show an extensive network of LAMA4 positive labelling on the tendon surface.

**Figure 3.**
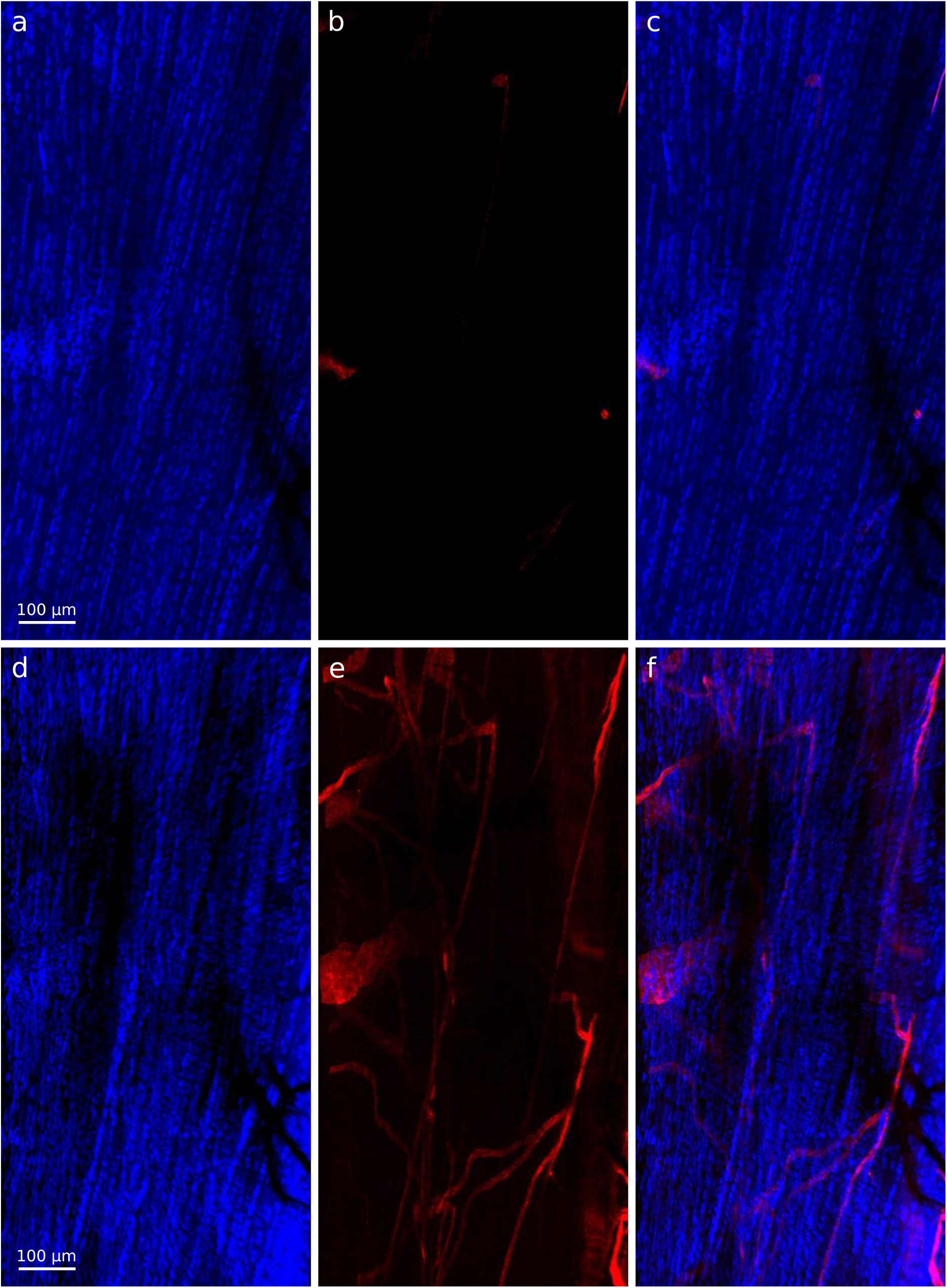
2D slices and maximum projection images of rat Achilles tendon core demonstrate information gained from 3D visualisation compared to 2D imaging. 2D slices (a-c) show sparse labelling for LAMA4 (b,c) within a highly cellular structure (a). Maximum intensity projections (d-f) of the same region shown in a-c demonstrate the presence of a complex network of LAMA4 positivity within the tendon core (e,f).

**Figure 4.**
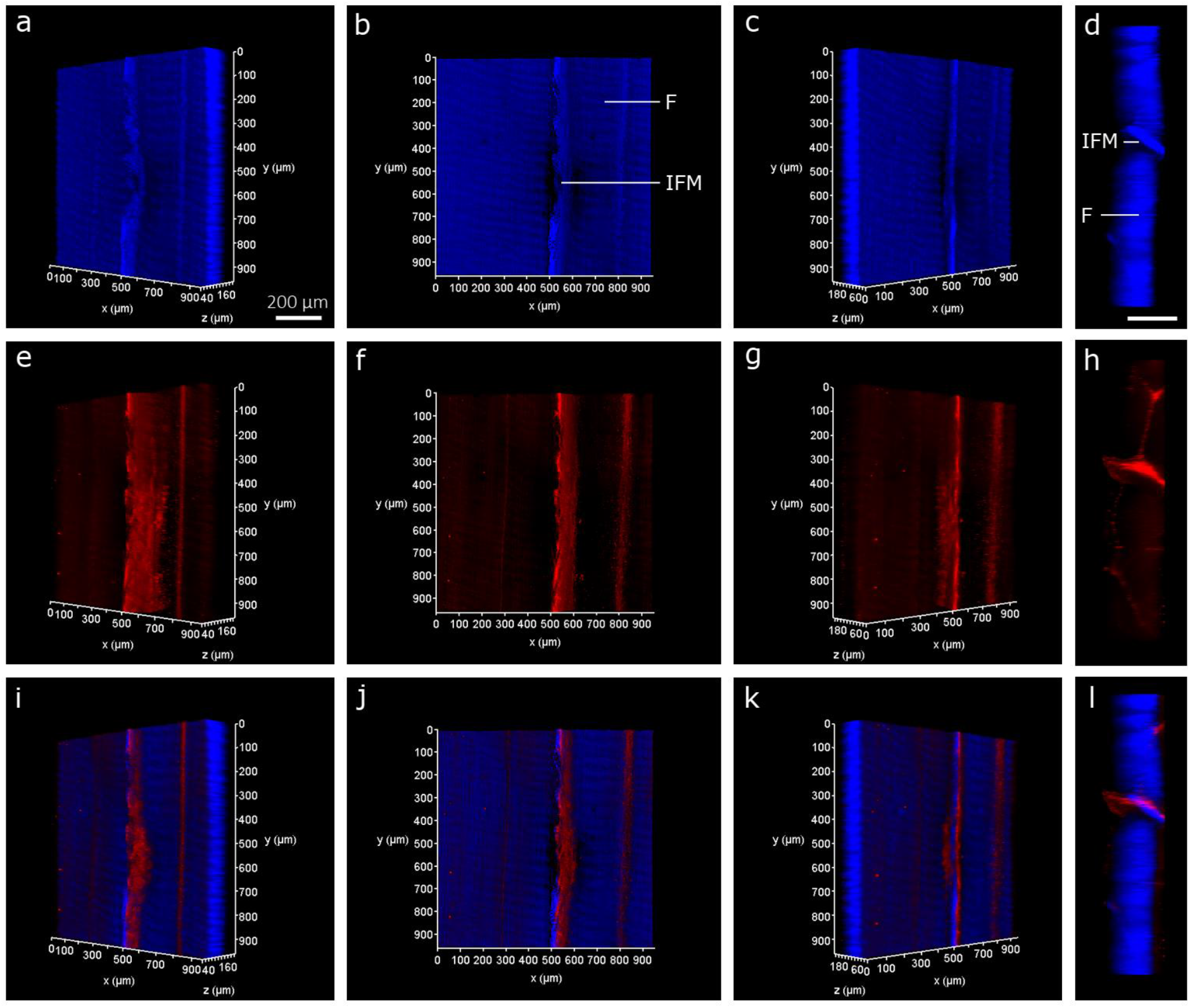
3D visualisation of LAMA4-immunolabelled equine superficial digital flexor tendon. 3D reconstructions of an equine superficial digital flexor tendon showing longitudinal nuclear (a-c) and LAMA4 (e-g) labelling, and with each channel overlaid to create a composite image (i-k) rotated at x=-45°, 0°, +45°. Transverse (xz) views of 3D reconstructions of equine SDFT (d,h,l) demonstrate signal present throughout the depth of the tissue. Interfascicular matrix (IFM) and fascicles (F) are denoted longitudinally (b) and transversely (d).

A key disadvantage to most commercially available optical clearing agents is the irreversibility of the clearing procedures. Here, we used an ethanol-based dehydration procedure for reverse clarification of Visikol HISTO™ solutions, followed by a previously described chemical drying process using HMDS to generate contrast for μ-CT [32]. Using grey-scale images captured from reconstructed μ-CT tomograms of reverse cleared and dried samples, we demonstrated that HMDS drying provides excellent contrast capable of visualising tendon surface and core structure in both species. Additionally, false colour 3D volume rendering provides indication of relative densities and enhances visualisation of tendon internal structure (**Figure 5 & Figure 6**).

**Figure 5.**
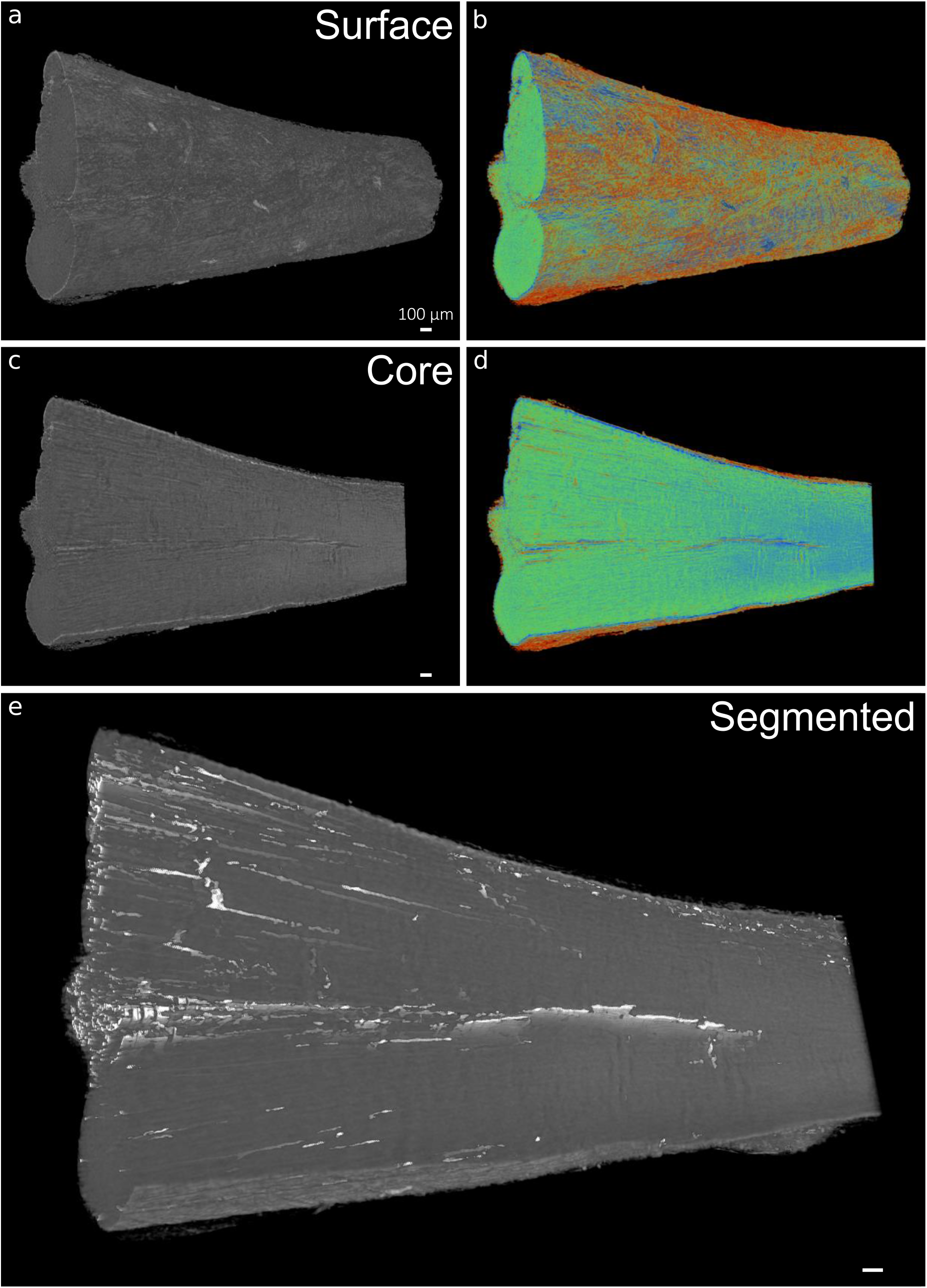
3D visualisation of x-ray micro-computed tomography-scanned rat Achilles tendon. 3D reconstructions in grey scale (a, c) and false-colour renderings (relative densities) (b,d) of rat Achilles tendon surface and core demonstrate the contrast between IFM and tendon substance created using HMDS drying. Automated segmentation allows 3D visualisation of IFM (white) in whole rat Achilles tendon (e).

**Figure 6.**
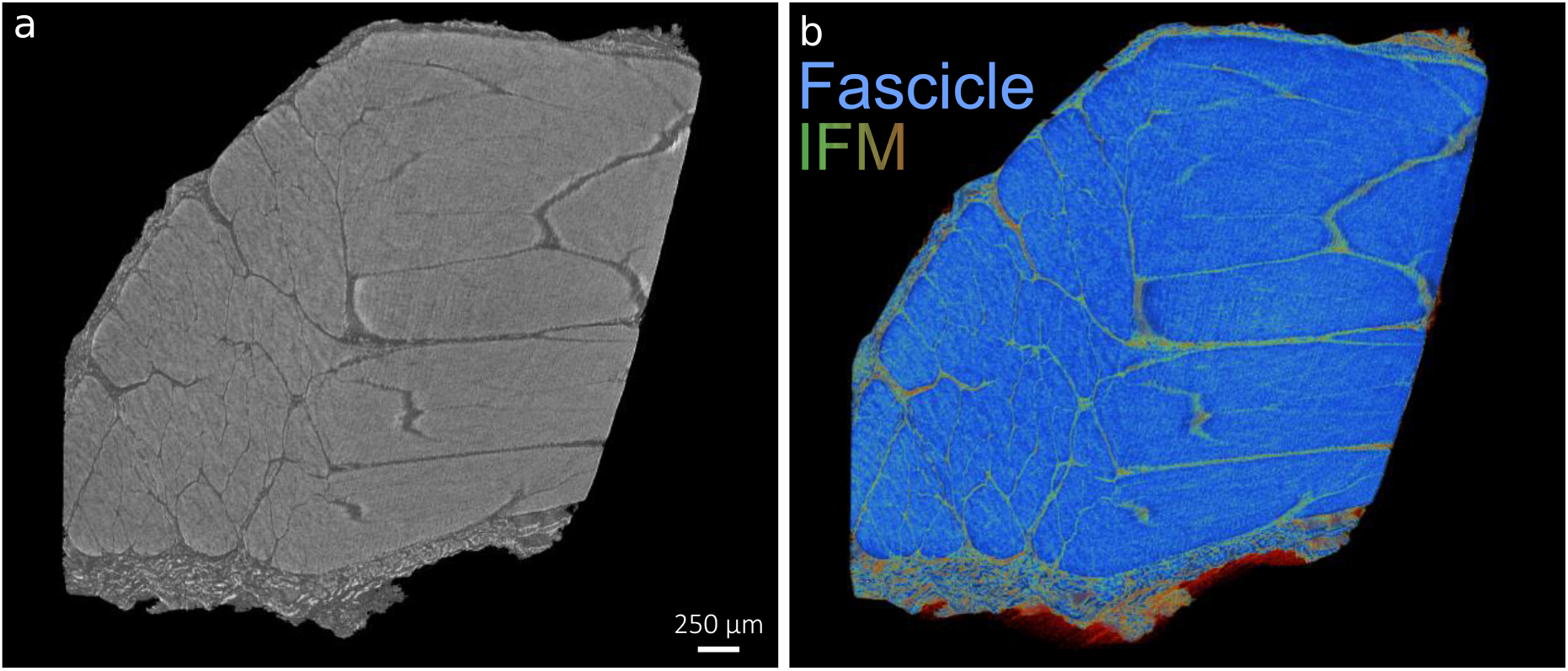
3D visualisation of x-ray micro-computed tomography scanned equine SDFT. 3D reconstructions in grey scale (a) and false-colour renderings (b) of equine SDFT demonstrate the contrast between IFM (green) and tendon fascicles (blue) using HMDS drying.

An important feature of these morphometric approaches is the scope they provide for detailed quantitative analysis of the tissue organisation. To demonstrate this utility, we therefore used an automated segmentation procedure to separate both IFM and fascicle structures in equine SDFT. By adapting pre-existing measurement modalities, designed for measurement of bone parameters (see **Table 1**), we were able to record tendon volume (TV), IFM volume relative to tendon volume (IV/TV), and IFM thickness (IFM.th) (**Figure 7**). To explore the possibility that such quantitative methods to assess tendon sub-structures may be impacted by artefacts introduced by optical clarification and reverse clearing, we scanned reverse cleared and non-cleared segments derived from the same tendon, that had both been HDMS-dried and compared IFM and tendon measurements. As expected, we found there was no difference in tendon volume as both samples (4.80-4.82 mm^3^) were analysed using the same volume-of-interest (1.75 mm × 1.75 × 2 mm). We found that IFM volume did not differ markedly between the HMDS only (0.26 mm^3^) and reverse cleared, HMDS-dried tendon samples (0.31 mm^3^). Hence, IFM volume to tendon volume was also similar between HMDS only (5.5%) and reverse cleared, HMDS-dried tendon (6.3%) in equine SDFT segments. IFM thickness measurements were also similar when comparing both samples, with reverse cleared tendon exhibiting a slightly thicker average IFM (0.022 mm) than HMDS only (0.019 mm), both values which fall within the range of IFM thickness (0.01-0.025 mm) measurements previously reported for an individual equine SDFT using 3D histological reconstructions [33].

**Table 1.** Description of 3D tendon morphometric parameters.

| **Original parameter name for bone analysis** | **Adapted tendon parameter name** | **Description of adaptation** | **Justification** | **Unit of measurement** |
| --- | --- | --- | --- | --- |
| Tissue volume (TV) | Tendon volume (TV) | Provides a total 3D volume measurement of scanned tendon (VOI). | Can be interpreted identically to tissue volume. | mm<sup>3</sup> |
| Bone volume (BV) | IFM volume (IV) | Provides a total volume measurement of binarised IFM within the VOI. | Can be interpreted identically to bone volume. | mm<sup>3</sup> |
| Percent bone volume (BV/TV) | Percent IFM volume (IV/TV) | Provides a proportion of the tendon volume occupied by IFM. | Can be interpreted identically to percent bone volume. | % |
| Trabecular thickness (Tb.Th) | IFM thickness (IFM.Th) | Provides a measurement of IFM thickness within the VOI. | Can be interpreted identically to trabecular thickness. | mm |

**Figure 7.**
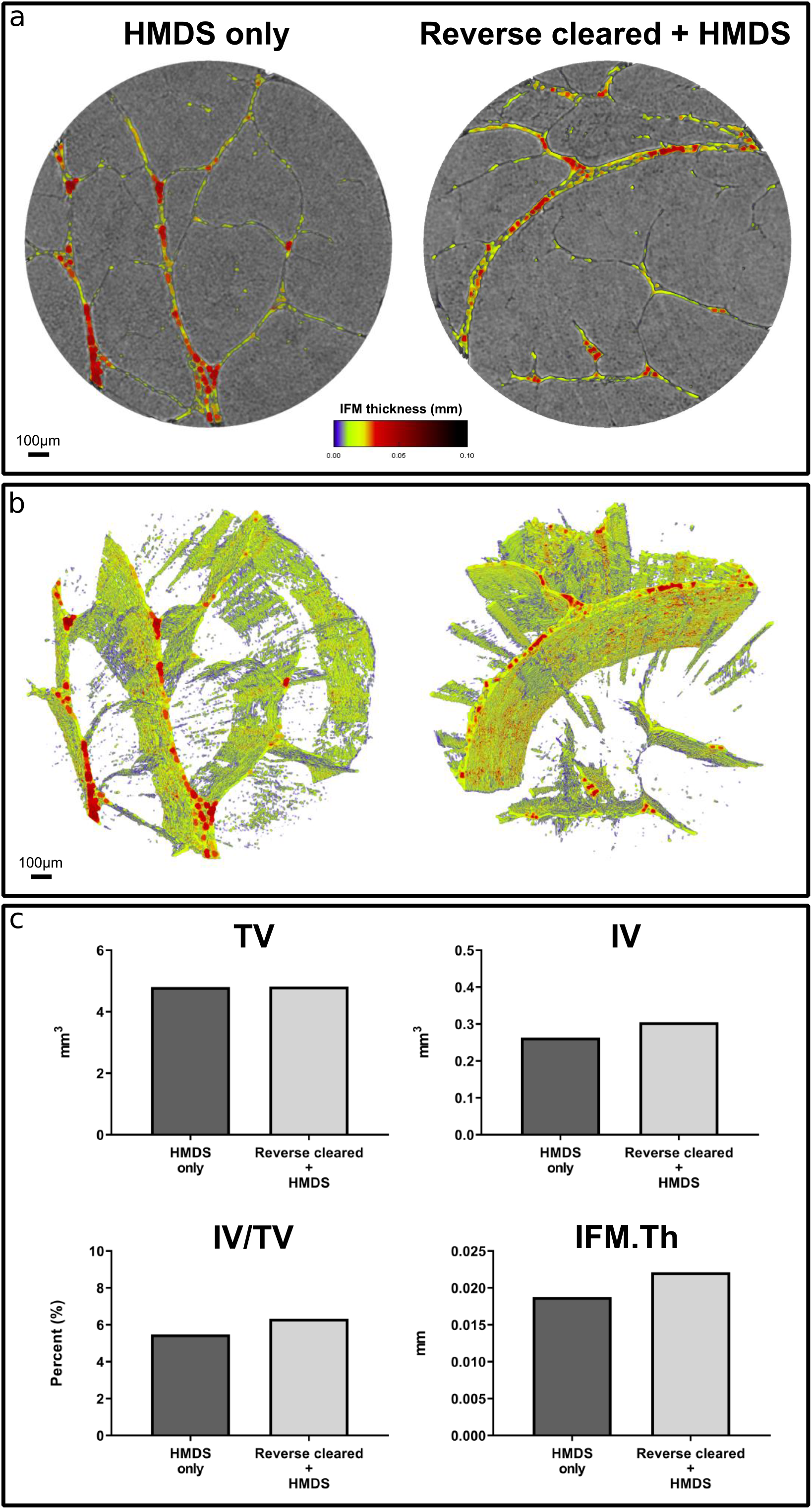
3D analysis of x-ray micro-computed tomography-scanned equine superficial digital flexor tendon. 3D reconstructions of segmented tendon and IFM thickness heatmaps (a-b) of chemically dried (HMDS only) and reverse cleared (Reverse cleared + HMDS) tendon. 3D analyses (c) demonstrate that both conditions share similar percentage of IFM volume (IV/TV) and IFM thickness (IFM.Th) measurements.

## DISCUSSION

We present a technique combining whole-mount confocal microscopy and μ-CT for the imaging of adult tendon, using a reversible clearing approach that renders tissue transparent for confocal imaging, followed by reverse clarification and chemical drying that permits contrast-enhanced μ-CT. We have demonstrated that not only does our methodology provide independent fluorescence and x-ray-based imaging modalities for dense connective tissues in the adult, but also enables both modalities to be combined for bimodal imaging of a single sample. This approach overcomes the limitations of traditional disruptive serial sectioning and en bloc staining procedures, introducing methods for visualising tissue structure within a single sample at a meso- and microscopic scale to elucidate the relationship between cell and ECM in the context of tissue structure and function.

We applied our bimodal procedure to both rat and equine tendons to demonstrate our method can be applied to small and large animal models. Hence, bimodal imaging offers the potential to image tendon structures for comparative biology purposes as well as for pathological conditions. Here, we used a novel marker of tendon basement membrane, LAMA4, to visualise equine and rat tendon sub-structure. LAMA4 is a component of laminin-8, laminin-9, and laminin-14, and has been described as a key regulator of basement membrane remodelling and function in adipose, cardiac, endothelial, hematopoietic, muscular, neuronal, and renal cell niches [34–37]. Our 3D imaging modality identified a complex network of LAMA4 immunolabelling in the core of tendons from both small and large animals, where it appears to localise predominantly to the IFM. This complexity can only be appreciated when viewing 3D reconstructions rather than 2D slices. Future studies could use our approach to investigate basement membrane remodelling and cell response to inflammation or injury in tendons and other soft tissues.

Our protocols also improve upon existing μ-CT modalities for visualising tendon structure, providing the ability to segment both fascicles and IFM from 3D reconstructions. Previous studies using contrast agents have presented procedures to segment and identify some but not all fascicles, but tendon structure could not be fully resolved [20]. However, sample drying using the HMDS protocol presented here provided adequate contrast for segmentation of both fascicles and IFM in both species and allowed quantification of IFM volume and thickness in equine SDFT. By modifying a reverse clarification procedure and combining with a chemical drying process, we present a method that allows cleared tendons to be imaged using contrast-enhanced μ-CT, facilitated by HMDS desiccation. This provides a rapid, cost-effective, non-specialised approach for μ-CT scanning, providing excellent contrast of tendon structure in both large and small animal models. However, we were unable to measure IFM volume in the rat Achilles, as the smaller IFM thickness in this tendon meant that, during segmentation, we were unable to discern between IFM and noise created by HMDS drying. Despite this caveat, this is the first study to report such methods for IFM volumetric measurements using μ-CT, finding that IFM represented a small volume of tendon (5-6%). Using segments derived from equine SDFT, we have also shown that mean IFM thickness, as measured by μ-CT, was between 19-22 μm for both HMDS-dried and bimodally imaged samples. This is in agreement with a previous study that described IFM thickness in the equine SDFT as measuring between 10-25 μm depending tendon regions [33], although we could not distinguish between IFM surrounding fascicles and fascicle bundles.

Another benefit to our bimodal workflow is the potential reduction in sample, and therefore animal, numbers required. Our approach nullifies the need for two distinct samples for two separate analysis techniques, providing the potential to reduce the number of animals used for scientific research, in line with the 3Rs principle [38]. Our methodology can also be implemented into pre-existing equipment workflows for confocal microscopy and μ-CT, and can also integrate open-source options in the absence of specialist software, such as Icy software for 3D whole-mount fluorescence visualisation and ImageJ/BoneJ for tendon morphometric analyses [39, 40]. Given we have only utilised one clarification method, future studies could investigate the effects of other clearing agents on tendon clarification, however other studies have compared a range of clearing solutions reporting no or limited effects on clearing of skeletally mature tendons [12, 15]. As clearing agents can have a tissue-specific effect, there is a greater need for understanding how clarification agents impact tendon ultrastructure, with previous studies demonstrating the destructive effects of different clearing agents when imaging GFP-labelled intestine samples [41]. Here, we have not directly determined whether tendon ultrastructure was affected, but no visual differences in tendon macroscopic structure were apparent pre/post-clearing and after reverse clarification. Our Visikol-based procedure also provides flexibility, with multiple stopping points throughout the protocol for both short and long-term storage without deterioration prior to clearing.

To enhance our methodology, 3D image registration of confocal and μ-CT images could be used for localisation of cellular and structural features of soft tissues such as tendon, whilst also enabling the identification of alterations in tissue sub-structure, such as basement membranes, with ageing and injury models. By combining imaging modalities, our method could visualise and analyse spatiotemporal differences in 3D, and would allow identification of injury-induced changes remote from injury site, especially for large animal tissues [42]. These approaches have already been attempted in renal and cardiac injuries in adult mice, demonstrating that 3D approaches can inform regional differences in tissue repair [43, 44]. Hence, our proposed protocol could be scaled down for smaller samples, such as mouse Achilles tendon, to investigate regional differences in cell:ECM interactions in murine models of tendon injury using 3D whole-mount preparation. Furthermore, antibody detection and novel RNA *in situ* hybridisation techniques could be used in conjunction with 3D registration to establish 3D molecular maps of localised soft tissue markers as evidenced in the developing chick heart [45]. Future studies could also pursue other potentially useful imaging modalities that could inform soft tissue structure-function relationships. For example, reversed clarified tissues can be histologically prepared for quantitative polarised light microscopy (qPLM) to investigate the structural arrangement of collagen and other fibrous proteins [46–48]. Therefore, 3D visualisation, image registration and subsequent analyses utilising bimodal imaging would greatly improve evaluation of a wide range of pre-clinical connective tissue disease models, as well as clinical samples.

### LIMITATIONS

To the authors knowledge, whole-mount fluorescent imaging of tendon has only been described in mouse embryonic tendons using transgenic systems [49], which pose little difficulty for 3D imaging due to immature collagens/ECM, smaller size, and visualisation via GFP-labelling. However, no such method has been described for the clearing of fluorescently-immunolabelled adult tendons from larger species where transgenic systems may not be financially or technically feasible. Despite this, our method does have some limitations. Firstly, antibody penetration appeared to differ in tendons from different species, with penetration throughout the depth of the rat Achilles (approximately 800 microns) but limited to 200 microns in the equine SDFT potentially due to lower surface area to volume ratio reducing ability for antibody diffusion. Similar limitations have been reported in osteochondral tissues labelled for large molecular weight ECM proteins, which also reported working depth as an issue for deep musculoskeletal tissue imaging [50]. Despite this, our method significantly improves upon the 20 – 25 microns imaging depth achieved in the SDFT without clarification reported previously [51, 52]. We also note that scanning large regions of SDFT and whole rat Achilles was performed at low magnification due to working distance of the objective, such that is was not possible to fully resolve cellular structures.

Limitations relating to μ-CT are also present in our method, mostly relating to scanning resolution. Segmentation of smaller structures and IFM remains a challenge despite the high resolution used to scan samples in this study. We found scans at a resolution of 1.6 microns provided high enough resolution to identify tendon sub-structures in both species, although quantification was not possible in rat Achilles tendon. While we were able to visualise IFM in the rat Achilles, the segmentation process generated some noise that we could not distinguish from IFM due to the smaller IFM size in rat compared to equine tendon. Therefore, quantification was not performed, as specialised segmentation procedures will need to be developed to improve filtering of noise in these small samples. As for automatic segmentation of equine SDFT, a small amount of IFM appears to be lost as a result of smoothing and filtering of 2D slices. In our 3D analyses, IFM thickness is calculated as a single mean value, therefore overall distribution of IFM thickness could not be described. Finally, we observed a small amount of shrinkage in both rat and equine tissues, despite this, our measures of IFM thickness are consistent with previous reports, therefore our method provides a viable approach to visualise and measure tendon morphometry using μ-CT.

To overcome these issues in either species, future scans could utilise higher-resolution or synchrotron-sourced x-ray tomography that offer greater imaging power. We also found that scanning without filters and using 180° projections provided the best acquisition of IFM in reconstructed images with reasonable scan time. Despite the aforementioned limitations, our approach permits sub-4-hour scans using benchtop μ-CT for downstream 3D volumetric analyses. Future studies could explore the benefits of various filters on scanning power when imaging tendon, whilst improvements to segmentation could be made with specialised segmentation routines such as adaptive local thresholding or machine-learning based segmentation [53]. We also noted that, while HMDS desiccation eliminated external artefacts observed previously when scanning samples in air [54], we did observe some x-ray ring artefacts in the μ-CT scans. This is most likely a side effect of the drying process itself as well as not using a filter during scans, however these artefacts were easily removed by automated reconstruction, ring smoothing and segmentation and as such do not pose any problems during 3D visualisation or analysis.

## CONCLUSIONS AND PERSPECTIVES

We report a novel bimodal imaging approach that utilises non-destructive, reversible clarification for 3D confocal microscopy of tendon which can be coupled with μ-CT procedures for analysis of tendon morphology. This methodology is a valuable addition to the imaging tools currently available to investigate both cellular and molecular biology of soft tissues, particularly in studies of comparative tendon biology to establish conserved structures and cell niche composition. By establishing a clarification protocol that enables study of tendon and other dense connective tissues, we address the lack of cost- and time-effective 3D fluorescent modalities for adult tissues. The introduction of chemical drying as a novel μ-CT preparation to enhance contrast of tendon structure provides the capacity to visualise and quantify tendon sub-structural features. Furthermore, both modalities can be performed independently of each other depending on experimental need and hypotheses. In conclusion, our bimodal imaging approach provides the capacity to visualise and analyse soft tissue structures, an advance which is likely to assist in pre-clinical or clinical investigation of connective tissue structure and pathologies.

## MATERIALS AND METHODS

### Sample preparation

Achilles tendons were collected from female Wistar rats (age = 13 weeks, n = 2), euthanised for a related tendon injury study in which contralateral uninjured tendons were not required for analysis. Tendons were washed in Dulbecco’s phosphate buffered saline (DPBS), fixed in 4% paraformaldehyde for 24 hours at RT and stored in PBS with 0.05% sodium azide at 4°C until processing. Forelimb SDFT were collected from female horses (n = 2, age = 7 and 23 years) euthanised at a commercial abattoir for reasons unrelated to tendon injury. Pieces (approximately 5 mm × 5 mm × 2 mm) were isolated from the mid-metacarpal region of the tendon, washed briefly in DPBS supplemented with 1% (v/v) antibiotic-antimycotic solution and fixed in 4% paraformaldehyde/10% neutral buffered formalin at RT for 24 h. Samples were stored in 70% ethanol at 4°C until processing.

### Fluorescent immunolabelling and clarifying procedures

Prior to whole-mount preparations, serial sections of equine SDFT and rat Achilles were fluorescently labelled with LAMA4 to validate expression in 2D and optimise antibody concentrations (see **Supplementary Methods**). For preparation of immunolabelled samples, equine SDFT segments and whole rat Achilles were processed according to Visikol™ guidelines. All steps were performed with orbital agitation at 60 RPM. SDFT segments were washed twice for 12 h with tris buffered saline (TBS) at RT, and permeabilised sequentially in 50% (v/v) methanol:TBS, 80% (v/v) methanol:dH_2_O, and 100% methanol for 2 h at 4°C. Samples were washed sequentially for 40 minutes at 4 °C with 20% (v/v) dimethyl sulphoxide (DMSO):methanol, 80% (v/v) methanol:dH_2_O, 50% (v/v) methanol:TBS, TBS, and TBS supplemented with 0.2% (v/v) Triton X-100 (0.2% TBS-TX100). For rat Achilles tendons, permeabilisation was performed as described for equine SDFT, with the exception that DPBS was substituted for TBS in all steps. Use of either saline solution does not impact labelling of tissue.

Prior to blocking, samples were incubated with a pre-blocking penetration buffer containing 0.2% TBS-TX100, 0.3 M glycine, and 20% DMSO for 6 h at 37 °C. Equine SDFT segments were blocked for 80 h at 37 °C in 0.2% TBS-TX100 supplemented with 6% (v/v) goat and 6% (v/v) donkey serum and 10% (v/v) DMSO. For rat Achilles tendons, blocking was performed in 0.2% TBS-TX100 supplemented with 6% (v/v) goat serum and 10% (v/v) DMSO. Primary antibody incubations for rabbit anti-LAMA4 (1:200; STJ93891, St. John’s Laboratories) for equine SDFT and 1:100 for rat Achilles tendons were performed at 37 °C for 80 h in respective antibody buffer containing TBS supplemented with TWEEN20 (0.2% v/v; TBS-TWEEN20), 3% (v/v) goat serum, 3% (v/v) donkey serum, and 5% (v/v) DMSO. For rat Achilles tendon antibody incubations, donkey serum was omitted.

Samples were washed three times for 2 h in wash buffer containing TBS supplemented with 0.2% (v/v) TWEEN20. Secondary antibody incubation was performed with goat anti-rabbit Alexa Fluor® 594 (A11037, Fisher Scientific) diluted at 1:250 for equine SDFT and 1:500 for rat Achilles tendon in antibody buffer for 36 h for equine SDFT and 72 h for rat Achilles tendon at 37 °C. Samples were washed five times for at least five minutes with wash buffer, before an overnight wash in wash buffer supplemented with DAPI (1:2000) for cell nuclei counterstaining. Samples were dehydrated as described above with increasing concentrations of methanol. Two-step tissue clarification was performed by immersing samples in Visikol HISTO-1 for 36 h for equine SDFT and 72 h for rat Achilles, followed by immersion in HISTO-2 for at least 36 h at RT. Samples were stored in HISTO-2 at 4 °C prior to confocal imaging.

### Confocal imaging

Samples were immersed in HISTO-2 on a glass-bottom dish fitted with a silicone chamber for imaging. Serial optical sections (equine SDFT z-stack = 200 μm; rat Achilles tendon z-stack = 1000 - 1500 μm) were acquired using a Leica TCS SP8 laser scanning confocal microscope with a motorised stage. Images were acquired with a HC PL FLUOTAR 10×/0.32 dry objective lens at a resolution of 1,024 x1,024 px, pinhole size set to 1 Airy unit, frame average set to 1, line average set to 8, and electronic zoom set to 0.75. Sequential scans of samples (rat Achilles: approx. 7mm × 2mm × 1.5mm; equine SDFT: approx. 1mm × 1mm × 0.2mm) were captured using lasers emitting light at 405 (blue channel; DAPI) and 561 (red channel; Alexa Fluor 594) nm to detect fluorescent signal, with low laser power (< 10%), and scanning speed set to 600 Hz. 3D rendering, segmentation, and projections were performed and visualised using Leica LAS X software (version 3.5.5) within the 3D module. Figures were produced using Inkscape (version 0.92).

### Reverse clarification and chemical drying for X-ray micro-computed tomography

Reverse clearing of tissues post confocal imaging were performed by sequential washes for a minimum of 3 h in 30%, 50%, 70%, 80%, 90%, 96%, and 100% (v/v) ethanol. Samples were immersed in HMDS solution (SIGMA: 52619) for 3-6 h and air-dried at room temperature overnight prior to X-ray micro-computed tomography.

### X-ray micro-computed tomography (μ-CT)

HDMS-dried equine SDFT and rat Achilles samples were affixed to a brass holder using dental wax and scanned using a Skyscan 1172F (version 1.5, Skyscan, Kontich, Belgium) with X-ray source at 40 kV tube voltage and 250 μA tube current with 1815 ms exposure time and 1.6 μm voxel size. 180° scans were performed with no filter, frame averaging at 5, with a rotation step at 0.2° and random movement set to 20. Slice reconstruction was performed in NRecon (version 1.7.1.0). Grey scale and false-colour volume renderings were produced using CTVox (version 3.3.0, Bruker, Belgium).

### 3D analysis of tendon IFM

3D analysis of equine SDFT pieces were performed using a cylindrical volume-of-interest (VOI) of 1.75 mm × 1.75 mm × 2 mm on all samples to avoid regions damaged by gross dissection of tendon. To process tendon plugs for 3D analysis, reconstructed images were processed using CTAn (version 1.17.7.1). To analyse tendon structure, we adapted existing workflows for bone analysis in CTAn (**Table 1**), to measure the volume of scanned tendon (TV), IFM volume (IV), relative IFM volume to tendon volume (IV/TV), and IFM thickness (IFM.Th). For 3D colour mapping of IFM thickness, BV and Tb.th were adapted as measures of IFM volume and thickness. These measurements were then used for 3D reconstruction and colour coded in CTVox (version 3.3.0). Graphs were produced in GraphPad Prism (version 8.0.0).

## LIST OF ABBREVIATIONS

|  |  |
| --- | --- |
| **3D** | Three-dimensional |
| **μ-CT** | x-ray micro-computed tomography |
| **C** | Celsius |
| **DAPI** | 4',6-diamidino-2-phenylindole dihydrochloride |
| **dH<sub>2</sub>O** | Deionised water |
| **DMSO** | Dimethyl sulfoxide |
| **DPBS** | Dulbecco's phosphate-buffered saline |
| **ECM** | Extracellular matrix |
| **HMDS** | Hexamethyldisilazane |
| **IFM** | Interfascicular matrix |
| **LAMA4** | Laminin Subunit Alpha-4 |
| **ms** | Milliseconds |
| **RPM** | Revolutions per minute |
| **SDFT** | Superficial digital flexor tendon |
| **TBS** | Tris-buffered saline |
| **TBS-T** | Tris-buffered saline supplemented with tween20 |
| **TBS-TX100** | Tris-buffered saline supplemented with triton-x100 |
| **VOI** | Volume-of-interest |

## DECLARATIONS

### Ethics approval and consent to participate

Rat Achilles tendons were obtained as waste tissue from a related study which was approved by the Royal Veterinary College ethics committee, conducted under a Home Office licence and complied with the Animal (Scientific Procedures) Act (1986). Collection of equine tendon was approved prior to commencement of the project by the Royal Veterinary College Ethics and Welfare Committee (URN-2016-1627b). Tendons were sourced from horses euthanised for reasons other than tendon injury at a commercial equine abattoir as a by-product of agricultural industry. The Animal (Scientific Procedures) Act 1986, Schedule 2, does not define collection from these sources as a scientific procedure.

### Consent for publication

Not applicable.

### Availability of data and materials

All data generated during this study will be uploaded to the Biostudies repository.

### Authors’ contributions

NM and CTT conceived the study design, performed all experimental work, analysis, visualisation, and produced the manuscript. APH contributed to development and optimisation of confocal imaging procedures. MH conceived and co-performed experimental design relating to μ-CT scanning, segmentation, analysis, and visualisation. NM, MH, APH, AAP, CTT interpreted data, critically revised the article, and approved the final manuscript for submission.

## SUPPLEMENTARY METHODS

### 2D immunolabelling of equine SDFT and rat Achilles tendon to establish optimum antibody concentrations

Longitudinal cryosections were cut from one SDFT (10 μm; 6 year old horse) and one rat Achilles tendon (12 μm; 13 week old female Wistar) that had been embedded in OCT and snap-frozen in n-hexane cooled on dry ice. Sections were adhered to glass slides and stored at −80 °C prior to immunolabelling. Sections were thawed, fixed with 4% PFA for 10 minutes and washed with TBS. Blocking conditions were consistent with those described for cleared samples used in the main study. Sections were incubated overnight at 4 °C with rabbit anti-LAMA4 primary antibody at dilutions ranging from 1:100 to 1:500 to establish optimum concentrations. Sections were washed with TBS, incubated with goat anti-rabbit Alexa Fluor® 594 secondary antibody (1:500, 2h, RT), coverslipped with Prolong Gold antifade mountant with DAPI (Invitrogen™ P36941) and cured overnight. Negative controls were included in which the primary antibody was omitted. Sections were imaged using a Leica TCS SP8 laser scanning confocal microscope a HC PL FLUOTAR 10×/0.32 dry objective lens.

A primary antibody concentration of 1:200 for the equine SDFT, and 1:100 for the rat Achilles provided optimal detection (Supp. Fig. 1). No non-specific labelling was detected in negative controls.

**Supp. Fig. 1:**
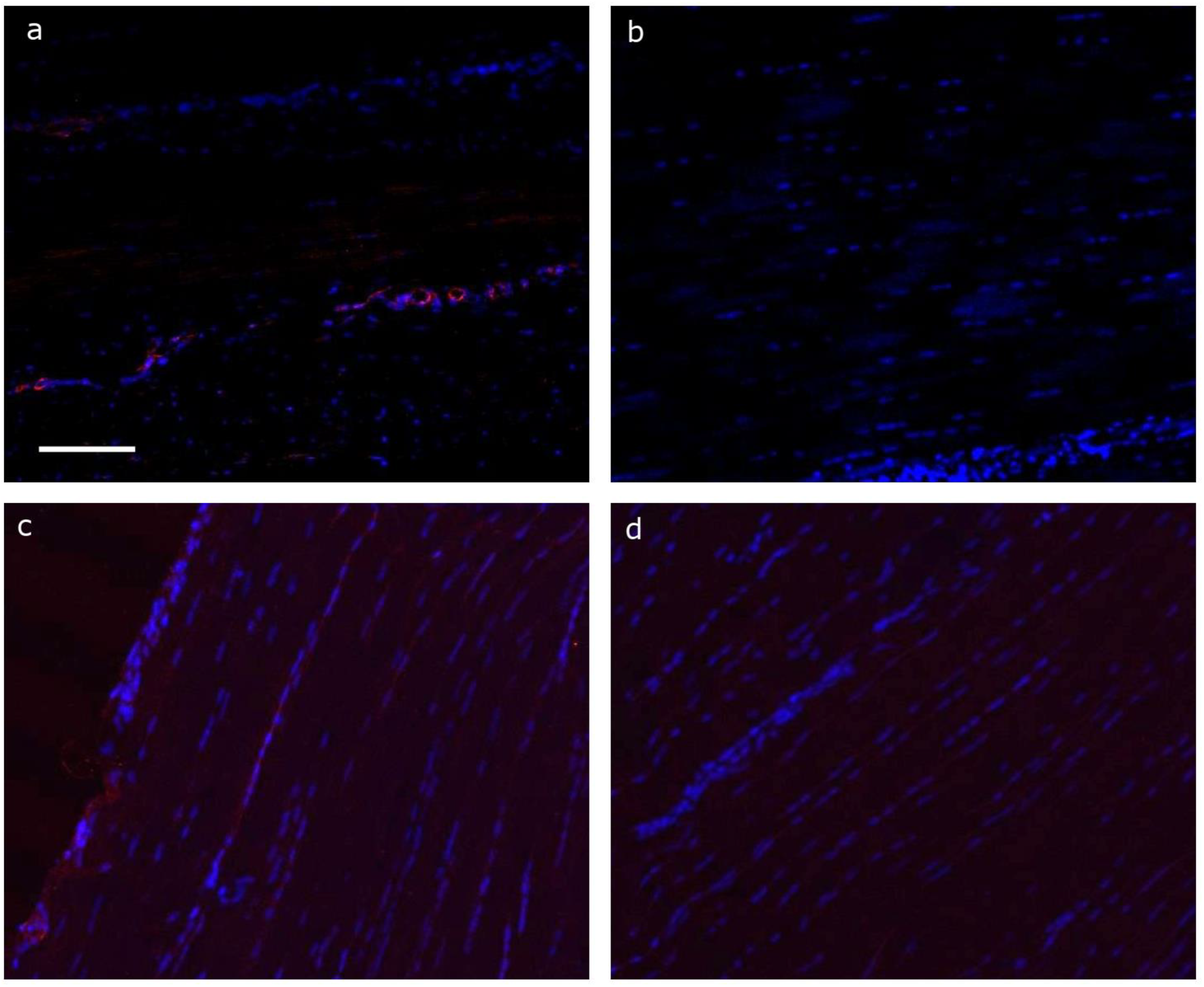
Representative 2D sections of equine superficial digital flexor and rat Achilles tendons, immunolabelled for LAMA4. Longitudinal sections were immunolabelled for LAMA4, using a primary antibody concentration of 1:200 in the equine SDFT (a) and 1:100 in the rat Achilles (c), detected with alexa-594 conjugated secondary antibody (1:500) and nuclei counterstained with DAPI. Negative controls, in which primary antibodies were omitted, were performed for both species (equine SDFT, b; rat Achilles, d). Scale bar = 100 μm.

